# Hip circumduction is not a compensation for reduced knee flexion angle during gait

**DOI:** 10.1101/520684

**Authors:** Tunc Akbas, Sunil Prajapati, David Ziemnicki, Poornima Tamma, Sarah Gross, James Sulzer

**Author notes:** Corresponding Author: Tunc Akbas, PhD, 204 E. Dean Keeton St, Austin, TX 78712.

## Abstract

It has long been held that hip abduction compensates for reduced swing-phase knee flexion angle, especially in those after stroke. However, there are other compensatory motions such as pelvic obliquity (hip hiking) that could also be used to facilitate foot clearance with greater energy efficiency. Our previous work suggested that hip abduction may not be a compensation for reduced knee flexion after stroke. Previous study applied robotic knee flexion assistance in people with post-stroke Stiff-Knee Gait (SKG) during pre-swing, finding increased abduction despite improved knee flexion and toe clearance. Thus, our hypothesis was that hip abduction is not a compensation for reduced knee flexion. We simulated the kinematics of post-stroke SKG on unimpaired individuals with three factors: a knee orthosis to reduce knee flexion, an ankle-foot orthosis commonly worn by those post-stroke, and matching gait speeds. We compared spatiotemporal measures and kinematics between experimental factors within healthy controls and with a previously recorded cohort of people with post-stroke SKG. We focused on frontal plane motions of hip and pelvis as possible compensatory mechanisms. We observed that regardless of gait speed, knee flexion restriction significantly increased pelvic obliquity (2.79°, p<0.01) compared to unrestricted walking (1.5°, p<0.01), but similar to post-stroke SKG (3.4°). However, those with post-stroke SKG had significantly greater hip abduction (8.2°) compared to unimpaired individuals with restricted knee flexion (4.2°, p<0.05). These results show that pelvic obliquity, not hip abduction, compensates for reduced knee flexion angle. Thus, other factors, possibly neural, facilitate exaggerated hip abduction observed in post-stroke SKG.

## Introduction

Stroke causes numerous impairments, including muscle weakness, spasticity, abnormal muscle coordination and altered proprioception, resulting in walking disorders. Stiff-knee gait (SKG), defined as reduced peak knee flexion angle during swing phase of the paretic side, is a common walking disorder following stroke (Deirdre Casey Kerrigan, Gronley, & Perry, 1991). Gait researchers have long assumed that people with post-stroke SKG compensate for reduced knee flexion with increased hip hiking, hip circumduction and/or vaulting (D Casey Kerrigan, Frates, Rogan, & Riley, 2000; Deirdre Casey Kerrigan et al., 1991; Perry & Burnfield, 1992). However, the causal relations between these compensations and reduced foot clearance have never been established. Stroke patients exhibit similar or higher foot clearance values compared to healthy individuals (Little, McGuirk, & Patten, 2014; Matsuda et al., 2017) suggesting one or more of these aforementioned compensations could be redundant. It is a commonly held belief that hip abduction, the main frontal plane component of hip circumduction, compensates for lack of knee flexion (Perry & Burnfield, 1992). However, the substantial energetic cost of hip abduction (Shorter, Wu, & Kuo, 2017) could make it the least desirable compensatory motion. Thus, in this work we investigated the necessity of hip abduction as a compensatory motion for reduced knee flexion.

Our previous work suggests excessive hip abduction may not be a compensation, but possibly the result of an abnormal coordination pattern emerging after stroke. We used a knee flexion torque assistance during pre-swing and observed an increased hip abduction angle during swing, instead of the expected reduction, in people with post-stroke SKG (Sulzer, Gordon, Dhaher, Peshkin, & Patton, 2010). The exaggerated hip abduction despite increased foot clearance suggested that abduction was not acting as a gait compensation. Other possible causes such as loss of balance, spasticity and reduced proprioception could not account for this phenomenon. Rather, excessive hip abduction could be part of a cross-planar abnormal coordination pattern, for example, reflex-based (Finley, Perreault, & Dhaher, 2008) or voluntary synergies (Cruz & Dhaher, 2008; Cruz, Lewek, & Dhaher, 2009; Neckel, Blonien, Nichols, & Hidler, 2008). Descriptive analyses of post-stroke gait have associated abnormal coordination with gait dysfunction. For instance, Clark et al. used non-negative matrix factorization to show that the number of coordination patterns in post-stroke negatively correlates with locomotor performance and clinical assessments compared to healthy individuals (Clark, Ting, Zajac, Neptune, & Kautz, 2010). Cases with fewer modules observed abductor activity coupled with sagittal plane muscles. Thus, accumulating evidence points to post-stroke hip abduction as part of abnormal coordination.

The concept of hip abduction in post-stroke SKG as an abnormal coordination pattern would be at odds with the widely accepted hypothesis that hip abduction is a compensation for reduced knee flexion (Perry & Burnfield, 1992). Here, we challenge the latter claim by determining how healthy individuals react to kinematically constrained knee flexion. If, as our earlier data suggest, abduction is not a compensatory motion, then we would expect healthy individuals to adapt to reduced knee flexion using other compensations, such as increased hip hiking (pelvic obliquity) or vaulting (increased plantarflexion of contralateral ankle during swing).

In this study, we simulated the mechanical constraints of SKG by restricting knee flexion in healthy individuals with an adjustable knee brace and observed the resulting compensations. Mechanical induction of gait deviations has been used to successfully quantify gait asymmetry (Shorter, Polk, Rosengren, & Hsiao-Wecksler, 2008) and evaluate energy expenditure (Hanada & Kerrigan, 2001). Lewek et al. mechanically induced SKG using a knee brace, finding that such a constraint results in higher metabolic cost (Lewek, Osborn, & Wutzke, 2012). Here we took a similar approach, but instead we examined how traditional compensatory parameters vary between those with post-stroke SKG and those with mechanically induced SKG. We additionally introduced other factors to more accurately simulate post-stroke gait, for example we matched walking speeds and added an ankle-foot orthosis commonly worn by individuals post-stroke. We predicted that more energy efficient compensations to reduce knee flexion, i.e. hip hiking, would facilitate foot clearance instead of abduction in healthy individuals. Differing reactions between post-stroke and restricted healthy individuals to similar knee motion would suggest that hip abduction is not a compensation for reduced knee flexion. This work distinguishes the impairment-related and compensatory joint motions in post-stroke gait, which will lead to improved clinical assessments and targeted therapy.

## Methods

Twelve unimpaired healthy individuals with no prior musculoskeletal injury gave written informed consent according to the guidelines approved by the University of Texas at Austin Institutional Review Board to participate in the experiment (Table S1).

The goal was to simulate the kinematic constraints of those with SKG in the unimpaired individuals and then compare with recorded data collected from participants with post-stroke SKG in previous study, where all participants were left-sided hemiparetics with knee range of motion at least 16° less on the effected side during swing phase (Sulzer et al., 2010). Since all patients had reduced knee flexion angle during swing, we restricted the knee with a commercial knee brace (Comfortland Medical, Mebane, NC) with a range-of-motion setting nominally at 0°. Since half of our patient sample wore an ankle-foot orthosis (AFO), we incorporated a commonly used AFO (Ossur, Reykjavík, Iceland) setting the ankle in a neutral, 90° ankle flexion position. Both orthoses were implemented on the left side to match the patient sample. We also further imitated our sample by matching gait speeds at 0.5 m/s. Thus, we used a 2×2×2 factorial design consisting of the factors of knee restriction, ankle restriction and walking speed. All subjects walked on a split-belt force treadmill (Bertec, Columbus, OH), which recorded ground reaction forces. Lower limb kinematic data were collected using an optical motion capture system (PhaseSpace Motion Capture, San Leandro, CA). Each of the three experimental factors consisted of two levels resulting in eight total conditions. Each healthy participant walked for three minutes for each condition, approximately 150 steps for slow walking speed and 200 steps for normal walking speed without receiving any prior practice. The *Restricted* condition consisted of both knee restriction and ankle restriction. The knee brace restriction was also implemented without the use of an AFO (*Brace*), and conversely the AFO was implemented without knee brace restriction (*AFO*). Lastly, subjects walked with no restriction at all while wearing the brace (*Free*). Each condition was implemented with slow (0.5 m/s) and normal (1 m/s) walking speeds, representing the walking speed of the post-stroke SKG cohort and typical comfortable healthy walking speed, respectively. The order of the conditions was randomized. Motion capture data was collected at 240 Hz and force measures from the instrumented treadmill were collected at 1 kHz. Figure 1 shows the representative overview of experimental setup demonstrating the *Restricted* condition. The data of nine individuals with post-stroke SKG collected from the baseline stage of previous study (Sulzer et al., 2010) was used to represent post-stroke SKG where participants walked at 0.5 m/s for two minutes.

**Figure 1:**
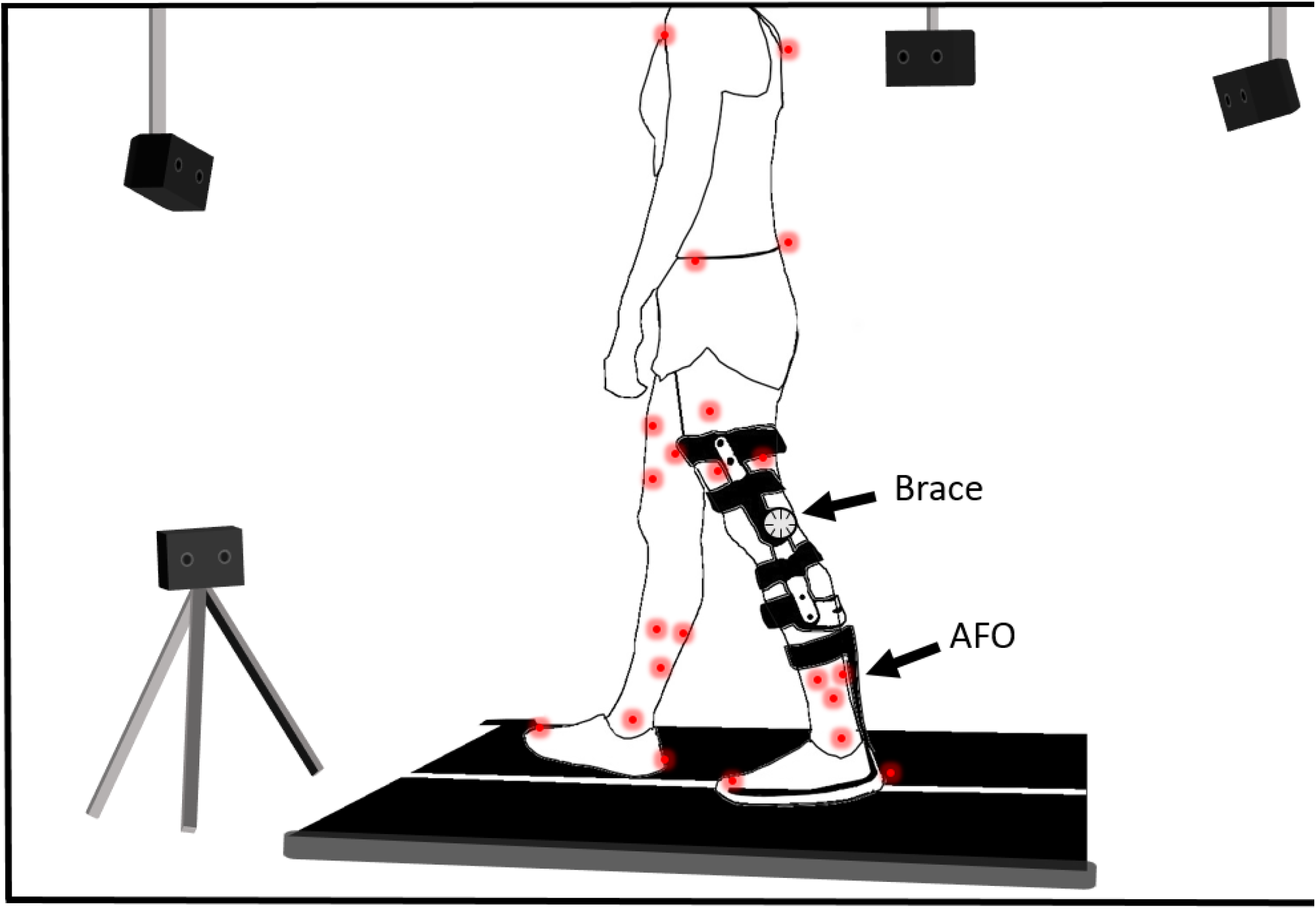
Schematic of experimental setup of mechanically induced SKG. A commercial medical knee brace and ankle foot orthosis (AFO) are used to imitate reduced knee flexion and orthotic assistance in post-stroke SKG, respectively. For the *Free* condition, the AFO and flexion angle constraint on the knee brace were removed, whereas for the *Brace* condition only the AFO was removed, and for the *AFO* condition only the flexion angle constraint was removed. Position of individual body segments was recorded with active LED markers (shown in red circles) via infrared cameras and gait events were detected using the force measures from the instrumented split-belt treadmill.

### Kinematic and Spatiotemporal Measures

Data was separated into gait cycles using left heel strikes for each participant corresponding to the given condition-speed pair. The heel-strike was detected using the instrumented split-belt treadmill based on a vertical force threshold of 10 N. The first 20 gait cycles were discarded to account for adaptation to the condition. Knee flexion, ankle plantarflexion and hip abduction angles of the ipsilateral (constrained/paretic) and contralateral (unconstrained/non-paretic) sides along with pelvic obliquity was extracted from a random selection of 25 gait cycles of each healthy participant for each condition to match the number of gait cycles collected from the individuals with post-stroke SKG from the previous experiment. We quantified hip circumduction as the hip abduction angle as opposed to the lateral displacement of malleolus (Lehmann, Condon, & Price, 1987) and coronal thigh angle (D Casey Kerrigan et al., 2000). The hip hiking is quantified by the coronal angle of the pelvis defined as pelvic obliquity (Michaud, Gard, & Childress, 2000). Range-of-motion (ROM) for each movement was defined as the difference between minimum and maximum joint angle measures in positive directions during pre-swing and swing phases of the gait cycle. The contralateral plantarflexion angle at toe-off was extracted to measure the amount of vaulting. Spatiotemporal characteristics were obtained including maximum toe height, maximum toe width, and toe height and width at minimum toe clearance from the ipsilateral (constrained/paretic) side (Figure 2). The maximum toe height was defined as the maximum vertical displacement of the toe marker. Maximum toe width was defined as the maximum lateral displacement of toe marker. Minimum toe clearance was quantified as the local minimum vertical displacement during swing phase (Winter, 1991). Finally, toe width at minimum toe clearance was quantified as the lateral displacement of toe marker at minimum toe clearance. Step asymmetry was quantified by the ratios of pre-swing times and swing times between ipsilateral (constrained/paretic) and contralateral (unconstrained/non-paretic) sides (Figure 2). Swing and pre-swing ratios were calculated by the ratio between swing phases of opposite limbs and the ratio between the durations in double support periods prior to swing of the corresponding limb respectively.

**Figure 2:**
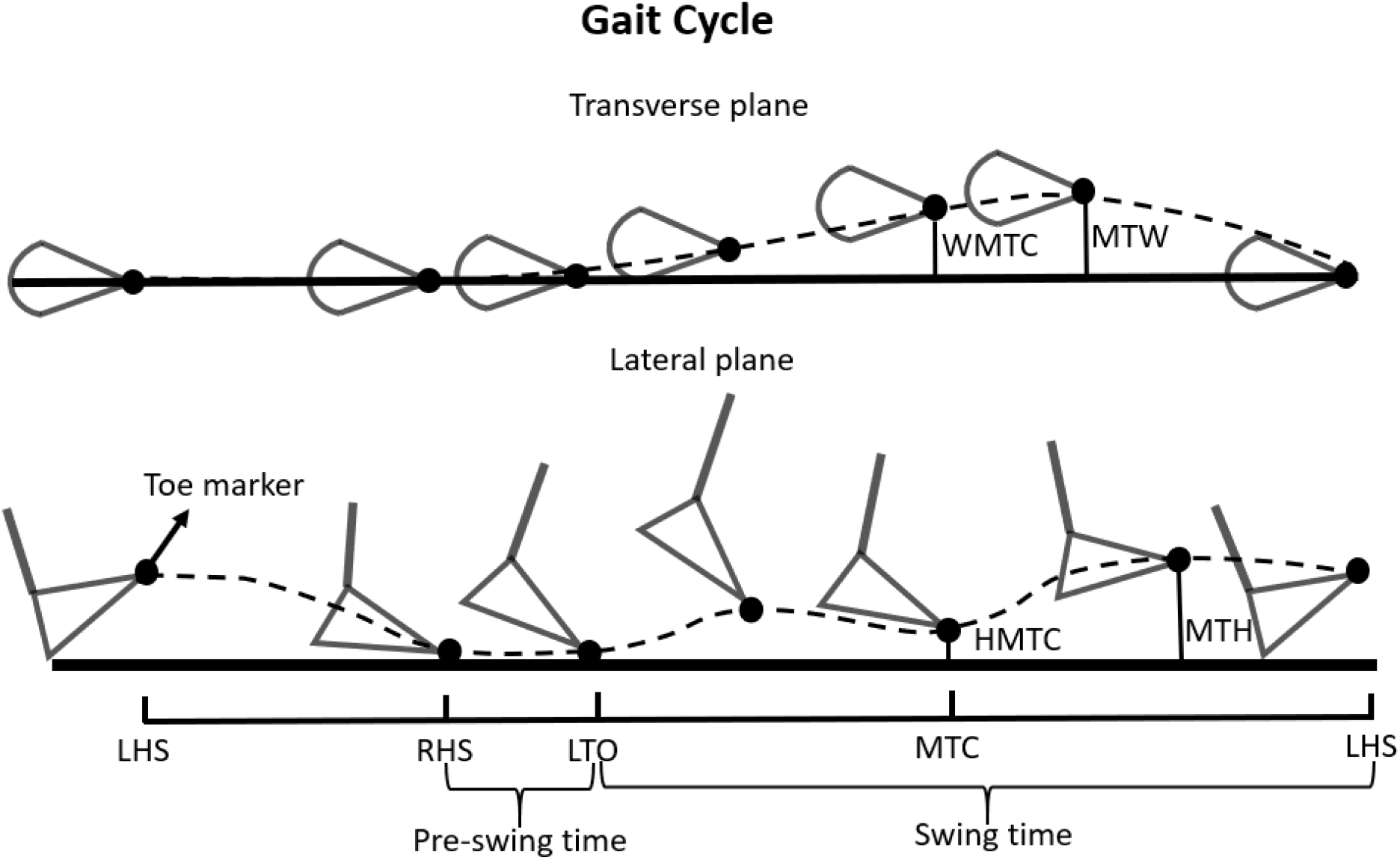
Selected spatiotemporal gait measures illustrated by left toe marker trajectory from lateral and transverse plane views. Maximum toe height (MTH), maximum toe width (MTW) and toe height (HMTC) and width (WMTC) measures at minimum toe clearance were obtained using the toe marker at the corresponding time instances during swing of the constrained/paretic side. The ratio of swing times (SR) and pre-swing times (PR) between constrained/paretic and unconstrained/non-paretic sides were used to measure gait asymmetry.

### Statistical Analysis

The collected data was analyzed using a linear mixed model (lme4) package (R Development Core Team, 2008). The first model included only the healthy participant pool. This model included the ROM measures from knee flexion, hip abduction and ankle plantarflexion of ipsilateral (constrained/paretic) side, and pelvic obliquity and spatiotemporal measures as dependent variables, with fixed effects of knee restriction, ankle restriction and walking speed. A linear mixed-effects model using the aforementioned factors, participant as a random effect and followed by Tukey-Kramer *post hoc* testing to evaluate the significance of the differences in the outcome variables between factors (α < 0.05).

The second model implemented the same linear mixed-effects model with the healthy participant pool at slow walking speed (0.5 m/s) with *Free* and *Brace* conditions as well as people with post-stroke SKG. We have accounted for repeated measures between *Free* and *Brace* conditions within the healthy group in this model using the same labels for the participants. Similar to the analysis in healthy individuals, we conducted Tukey-Kramer *post hoc* testing with the joint angle ROM measures and spatiotemporal measures as dependent variables. We examined differences between the participants with post-stroke SKG from the previously collected data and the two healthy conditions at matched speeds (*Free* and *Brace*).

We ran the Shapiro-Wilk normality test for all the outcome measures within corresponding factors for the first model and corresponding groups for the second model to confirm the normality of the data sets (*p*>0.05).

## Results

### Comparisons within healthy individuals

Average gait trajectories for all the aforementioned kinematic measures for all subjects in *Free, Brace, AFO and Restricted* conditions with slow and normal walking speeds are shown in Figure 3. A summary of the outcome measures for each condition can be found in Table S2. The following highlights the statistical comparisons based on the linear-mixed model.

**Figure 3:**
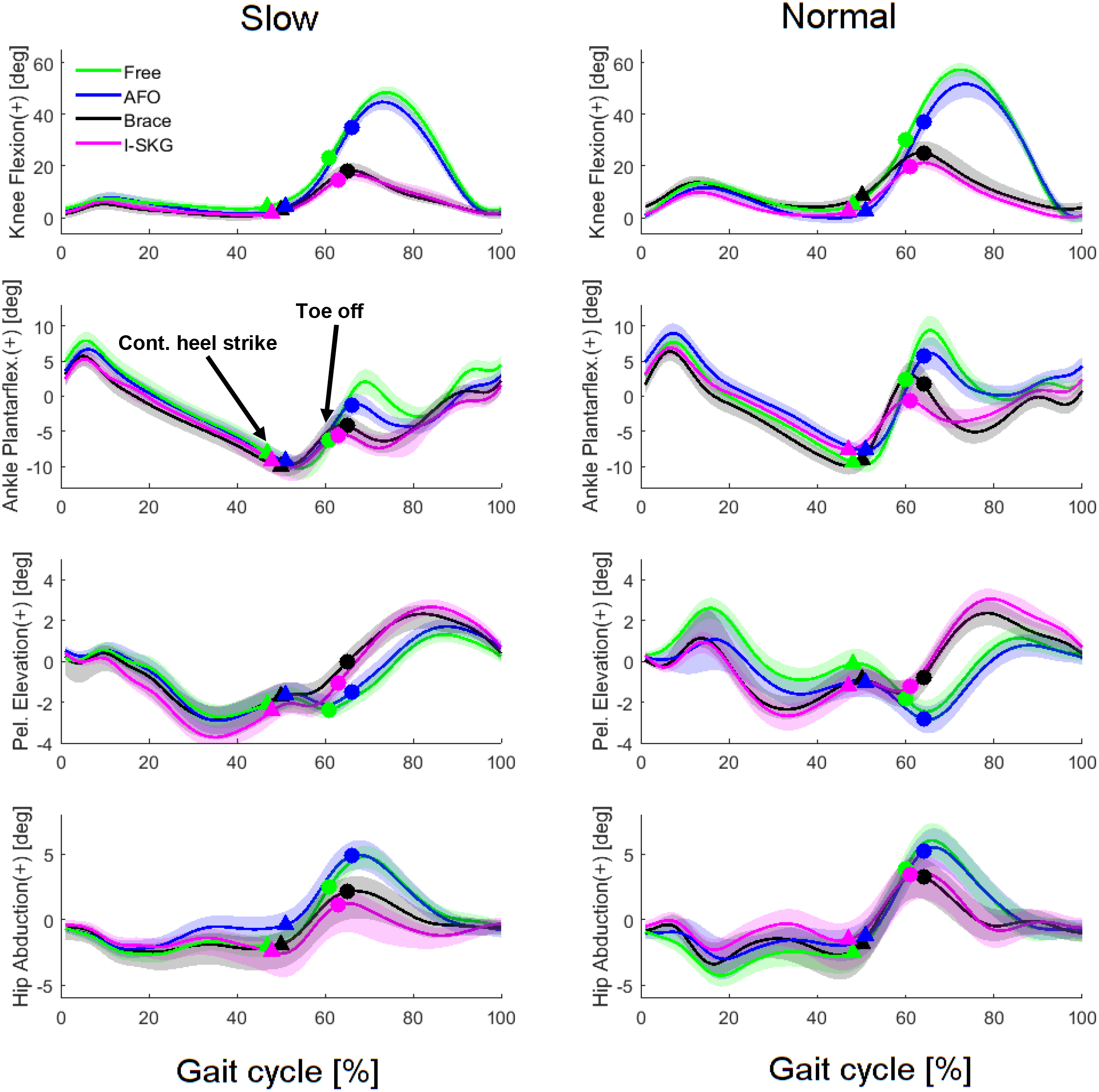
Healthy individuals compensate with increased pelvic obliquity and decreased ankle plantarflexion in response to reduced knee flexion. The knee flexion, ankle plantarflexion, pelvic obliquity and hip abduction angles for the constrained side under different conditions (*Free, AFO, Brace* and *Restricted*) are given for slow (left) and normal (right) walking speeds. The mean values and the standard errors are shown by solid lines and shaded areas, respectively. Contralateral heel strike is delineated by triangles and toe-off by circles. Regardless of walking speed, the introduction of knee restriction increased pelvic obliquity and decreased hip abduction in healthy controls.

We observed a main effect of knee restriction on knee flexion ROM (*F*_(1,76)_ = 158, *p*< .001) and ankle flexion ROM (*F*_(1,76)_ = 16.4, *p* < .001), both reduced significantly in *Brace* condition (*t* = −12.6, *p* < .001, *t* = 4.35, *p* < .001, respectively) compared to *Free* condition. Knee restriction also affected pelvic obliquity (*F*_(1,76)_ = 17.1, *p* < .001) but there was no significant difference in hip abduction (*F*_(1,76)_ = 2.27, *p* = .136). For instance, pelvic obliquity ROM increased in *Brace* condition compared to the *Free* condition (*t* = 5.26, *p* < .001). We observed the main effect of ankle restriction on ankle plantarflexion ROM (*F*_(1,76)_ = 6.00, *p* = .017), significantly reduced from the *Free* condition to the *AFO* condition (*t* = −2.61, p = .006). As expected, ankle restriction did not affect compensatory parameters such as hip abduction (*F*_(1,76)_ = 0.01, *p* = .908) or pelvic obliquity (*F*_(1,76)_ = 0.41, *p* = .53). Vaulting was not affected by knee restriction (*F*_(1,76)_ = 0.05, *p* = .824) or ankle restriction (*F*_(1,76)_ = 0.14, *p* = .708). When comparing the joint angle ROM measures between the *Brace* and *Restricted* conditions, i.e. the interactive effect of the ankle and knee restriction, there was no significant interaction effect for any of the measures (*p* > .05).

As expected, walking speed affected multiple kinematic variables, including knee flexion ROM (*F*_(1,76)_ = 9.46, *p* = .003) and ankle flexion (*F*_(1,76)_ = 5.07, *p* = .042), both decreased significantly in slow walking speed (*t* = −3.07, *p* = .003, *t* = −2.06, *p* <. 042, respectively). However, we did not observe any significant interactive effects between the walking speed and restrictions on knee and ankle (p > .05). That is, the change in speed did not modulate the relations between kinematic variables reported above.

Maximum step height was significantly affected by knee restriction (*F*_(1,76)_ = 15.94, *p*< .001) and toe height at minimum toe clearance (HMTC) was significantly affected by ankle restriction (*F*_(1,76)_ = 6.91, *p*< .010). The maximum step height was significantly decreased in *Brace* condition (*t* = −3.84, *p* < .001) compared to the *Free* condition, whereas HMTC was significantly increased in *AFO* (*t* = 2.41, *p* < .018) condition compared to *Free* condition. On the other hand, the maximum step width was significantly affected by knee restriction (*F*_(1,76)_ =6.25, p = .015) whereas no significant effect was observed in ankle restriction (*F*_(1,76)_ = 0.01, *p*= .918). The maximum step width was increased in the *Brace* condition (*t* = 2.45, *p* = .017) compared to the *Free* condition. In terms of gait symmetry, the swing-time ratio was only affected by knee restriction (*F*_(1,76)_ =14.94, *p*< .001) and pre-swing time ratio was only affected by ankle restriction (*F*_(1,1959)_ =5.68, *p*= .012). The swing time ratio was significantly increased in the *Brace* condition (*t* = 3.86, *p* < .001) and the pre-swing time ratio was significantly decreased in the *AFO condition* (*t* =−2.38, *p* = .020) compared to the *Free* condition.

As expected, walking speed affected multiple spatiotemporal measures, including maximum step height (*F*_(1,76)_ = 14.5, *p* < .001) and HMTC (*F*_(1,76)_ = 10.1, *p* = .002), both decreasing significantly in slow walking speed (*t* = −3.78, *p* = .003, *t* = −3.22, *p* <. 002, respectively). We did not observe any significant interactive effects between the walking speed and restrictions on knee and ankle (*p* > .05) for spatiotemporal measures, similar to the interactive effects between walking speed and knee and ankle joint ROM measures.

### Comparisons to post-stroke SKG

We compared the kinematic and spatiotemporal outcomes between the *Brace* and *Free* walking conditions at slow walking speeds and the post-stroke SKG group walking at the same speed. Our analysis only focused on the *Brace* condition because it is a more well-controlled condition compared to both ankle and knee restriction. Furthermore, *post-hoc* analysis did not indicate any significant differences between the *Brace* and *Restricted* conditions for kinematic and spatiotemporal outcome measures. Gait trajectories are illustrated in Figure 4 and corresponding ROM measures were shown in Supplementary Figure 1 and statistical comparisons between groups are shown in Table 1. The following is a summary of the statistical comparisons in selected parameters.

**Figure 4:**
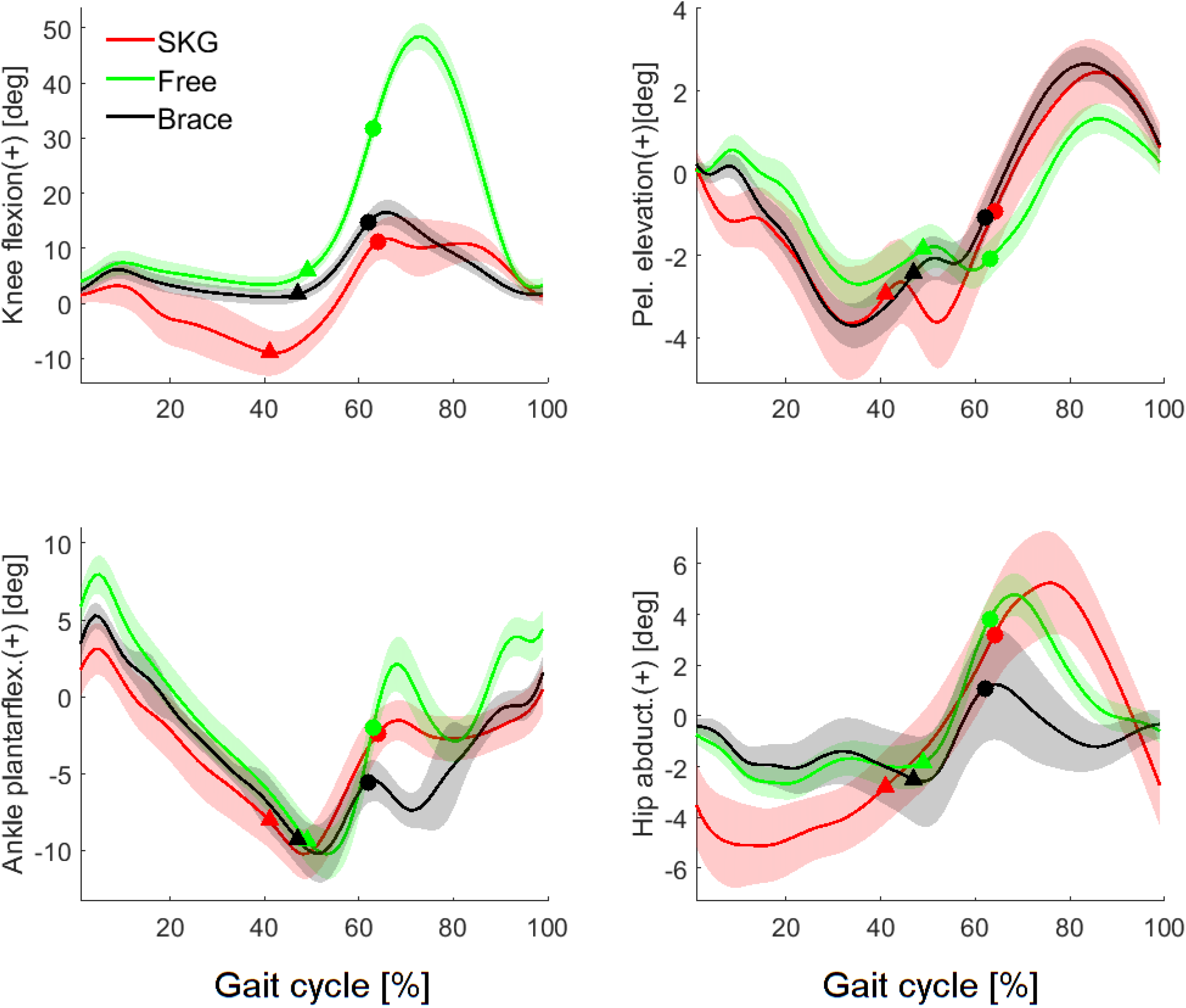
Increased hip abduction angle in stroke contrasts with decreased abduction in the *Brace* condition. The knee flexion, hip abduction and ankle flexion angles for the constrained/paretic side and pelvic obliquity for healthy individuals with *Free* and *Brace* conditions speed-matched with people with post-stroke SKG. The standard errors are shown by shaded areas. Contralateral heel strike is delineated by triangles and toe-off by circles. Note that despite similar knee flexion between the *Brace* condition and *SKG* group, the *Brace* condition resulted in reduced hip abduction while the *SKG* group had increased hip abduction.

**Table 1:**
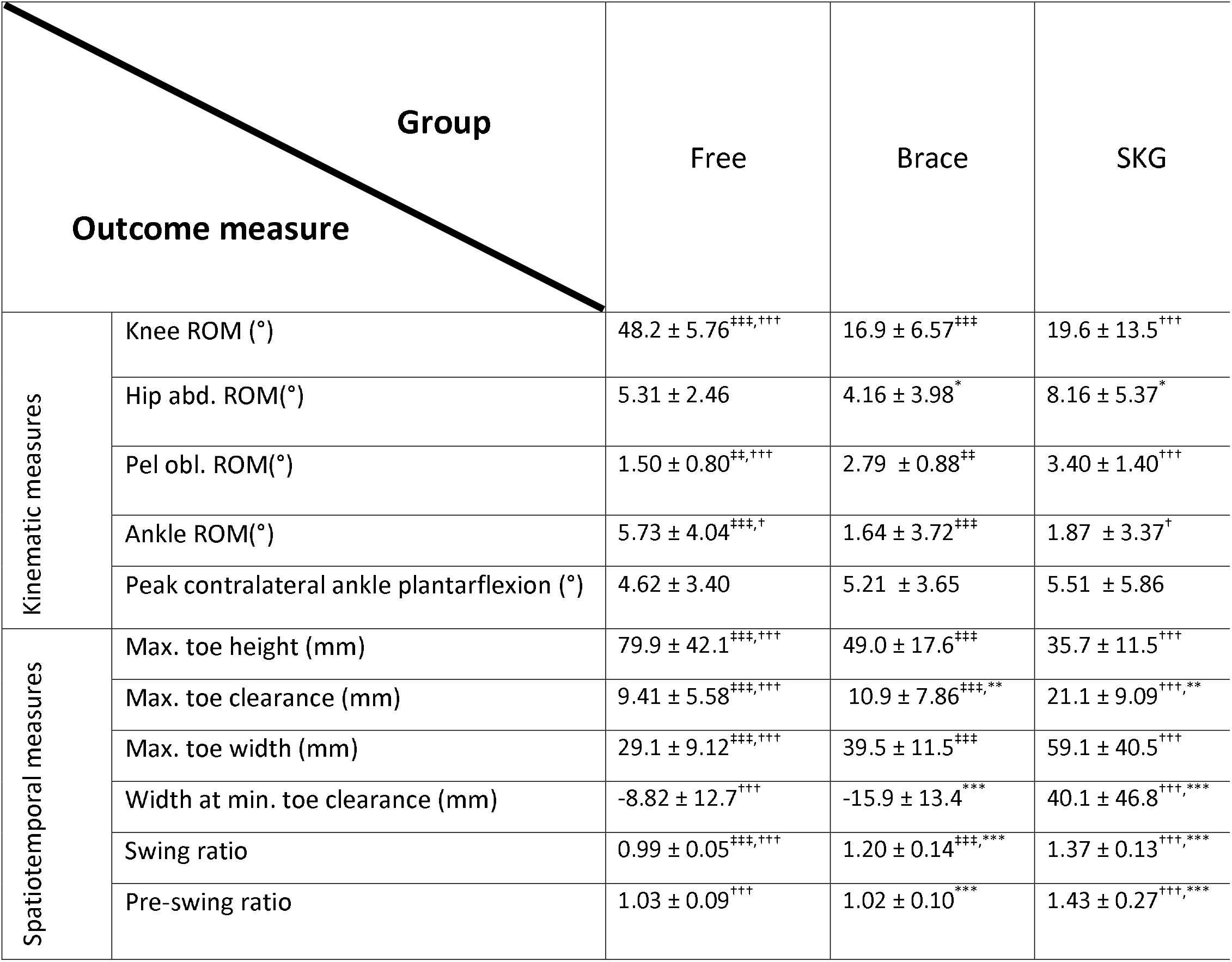
Summary of the joint angle ROM and spatiotemporal gait measures for healthy gait with and without knee brace restriction and post-stroke SKG. The measures indicate the mean values and standard deviations. The level of significance between *SKG* and *Brace* condition is denoted as * *p* < 0.05, ** *p* < 0.01, *** *p* < 0.001; between *Brace* and *Free* conditions denoted as ‡ *p* < 0.05, ‡‡ *p* < 0.01, ‡‡‡ *p* < 0.001; and between *Free* and SKG conditions denoted as † *p* < 0.05, †† *p* < 0.01, ††† *p* < 0.001.

The knee flexion ROM in the *Brace* condition and *SKG* were not significantly different (*t* = 0.73, p = .467) but the *SKG* group was significantly lower than the *Free* condition (*t* = 8.88, *p* = .001). Similarly, there were no significant differences in ankle plantarflexion ROM between the *Brace* condition and *SKG* (*t* = 0.26, *p* = .788). However, the *Free* condition had significantly higher ankle plantarflexion compared to the SKG group (*t* = 2.78, *p* < .013). There was no significant difference in pelvic obliquity ROM between the *Brace* condition and *SKG* group (*t*=3.02, *p* = .103) but the *SKG* group was significantly higher than the *Free* condition (*t* = 5.25, *p* <.001). In addition, hip abduction in post-stroke SKG was significantly higher than the *Brace* condition (*t*= 2.52, *p* = .012). There was no significant difference in vaulting between the *SKG* group and *Brace* condition (*t* = 0.53, *p* = .585) or between the *SKG* group and *Free* condition (*t*= 0.93, *p* = .351).

Spatiotemporal characteristics differed between groups. The maximum toe height was significantly higher for the *Free* condition compared to the *SKG* (*t*= 4.99, *p* < .001). Maximum toe width in the *Free* condition was significantly lower than the *SKG* group (*t*= 3.13, *p* < .001). The toe height at minimum toe clearance in the *SKG* group was significantly higher than both the *Free* (*t*= 4.08, *p* < .001) and *Brace* conditions (*t*= 3.49, *p* < .001). Toe width measures at minimum toe clearance in *SKG* was significantly higher compared to the *Free* (*t*= 3.79, *p* < .001) and *Brace* (*t*= 4.38, *p* < .001) conditions.

The pre-swing time ratio in *SKG* group was significantly higher compared to *Free* (*t*= 5.74, *p* <.001) and *Brace* conditions (*t*= 5.34, *p* < .001). Similarly, swing ratio in *SKG* group was significantly higher than the *Brace* condition (*t*= 7.57, *p* < .001) and the *Free* condition (*t*= 3.92, *p*< .001).

## Discussion

Hip abduction has long been assumed to be a compensation for reduced knee flexion angle during swing, and accordingly, excessive abduction in post-stroke SKG has been attributed to the same mechanism (Perry & Burnfield, 1992). To test this hypothesis, we applied kinematic constraints at the knee and ankle in healthy gait to imitate the sagittal plane kinematics of post-stroke SKG and evaluated resulting compensations. In response to reduced knee flexion, healthy individuals compensated with greater pelvic obliquity with no significant change in hip abduction, regardless of gait speed. Individuals with post-stroke SKG exhibited a similar increase in pelvic obliquity, but in contrast to healthy individuals with constrained knee flexion, higher hip abduction was observed. In addition, the minimum toe clearance of people with post-stroke SKG was also higher than those with mechanically induced SKG, indicating that the excessive abduction was not necessary for foot clearance. In summary, our results show that hip abduction is not a gait compensation for reduced knee flexion angle.

Our findings suggest that excessive hip abduction is unnecessary to facilitate swing phase toe clearance in those after post-stroke SKG, strongly questioning its use as a compensatory motion. Stroke participants exhibited the same sagittal plane joint ROM as healthy individuals with restricted knee motion and also exhibited the same compensatory motions indicated by the pelvic obliquity ROM. However, in stark contrast to healthy individuals with mechanically reduced knee flexion, those with post-stroke SKG walked with substantially higher hip abduction and toe clearance, seemingly with no biomechanical benefit. Previous work has reported excessive contributions of the ankle plantarflexor (soleus) and abductor muscles (gluteus medius) during forward propulsion and swing phase of the paretic side during post-stroke gait, indicating abnormal coordination similar to what we hypothesized (Hall, Peterson, Kautz, & Neptune, 2011). Our own work revealed that increased knee flexion angle and toe clearance provided by exoskeletal assistance resulted in greater hip abduction in post-stroke SKG which could not be accounted by biomechanical factors (Sulzer et al., 2010). Further analysis suggested that a cross-planar reflex coupling initiated by spastic rectus femoris co-activated with gluteus medius (Akbas, Neptune, & Sulzer, In review). Thus while excessive hip abduction should be expected to avoid excessive ankle plantarflexion, for example, during equinus deformity of the foot (Kinsella & Moran, 2008), the cause of excessive abduction in those with only reduced knee flexion could be due to non-biomechanical causes such as abnormal coordination (Brunnström, 1970).

Instead of abduction, we found increased pelvic obliquity as the primary compensation for reduced knee flexion. Earlier work has shown only increased pelvic obliquity during toe-off in post-stroke gait compared to healthy gait, without any significant changes in hip abduction (Cruz et al., 2009; Matsuda et al., 2016). Further, in those with post-stroke hemiparesis, pelvic elevation and knee flexion were inversely correlated with the walking speed, whereas no correlation was observed with hip abduction (Stanhope, Knarr, Reisman, & Higginson, 2014). Not only does this evidence suggest that pelvic obliquity is the primary compensation for reduced foot clearance, but additional work indicates the cost of abduction (Shorter et al., 2017). Increased circumduction magnitude is exponentially correlated with the cost of metabolic energy during walking (Shorter et al., 2017). Our results in healthy individuals conclusively add to this literature by directly illustrating that pelvic obliquity is the primary compensation for reduced knee flexion, whereas hip abduction is not a compensation.

There were additional differences between post-stroke and healthy individuals. For instance, higher temporal asymmetry was observed in post-stroke SKG compared to healthy gait with restricted knee motion. This difference can be explained by the increased swing time due to excessive abduction. Increased minimum toe clearance in post-stroke SKG compared to those with mechanically restricted knee motion indicates an overcompensation following toe-clearance. This overcompensation could be due to the lack of proprioceptive feedback following post-stroke hemiparesis (Keenan, Perry, & Jordan, 1983). Alternatively, the increased toe height at minimum toe clearance could be due to ankle impairments (Basmajian, Kukulka, Narayan, & Takebe, 1975; Olney, Griffin, Monga, & McBride, 1991).

There are limitations to this study that prevent greater generalizations. For instance, we cannot make conclusions regarding other kinematic abnormalities such as foot drop or the knee hyperextension prior to swing observed in people with post-stroke SKG. Knee hyperextension (genu recurvatum) and foot drop, while common in SKG, will not increase effective limb length and thus is a very unlikely contributor to supposed compensatory abduction. Our comparison was limited to kinematic and spatiotemporal factors and did not simulate motor control or proprioceptive contributions to gait compensations. While it is feasible that proprioceptive loss could contribute to increased hip abduction in post-stroke SKG, it is unlikely this would result in such an energetically costly compensation over a long period of time.

In conclusion, our data shows that hip abduction is not a necessary compensation for reduced knee flexion during gait, in direct opposition to widely held beliefs of abduction’s compensatory role. Instead, pelvic obliquity is the primary compensatory motion associated with reduced swing phase knee flexion ROM. <u>Together with previous findings, these data suggest that excessive hip abduction in those with post-stroke SKG could originate from a non-biomechanical cause, such as an abnormal coordination pattern. The correct characterization of compensation and impairment will lead towards improved treatment strategies and interventions</u>.

---

**Figure S1:**
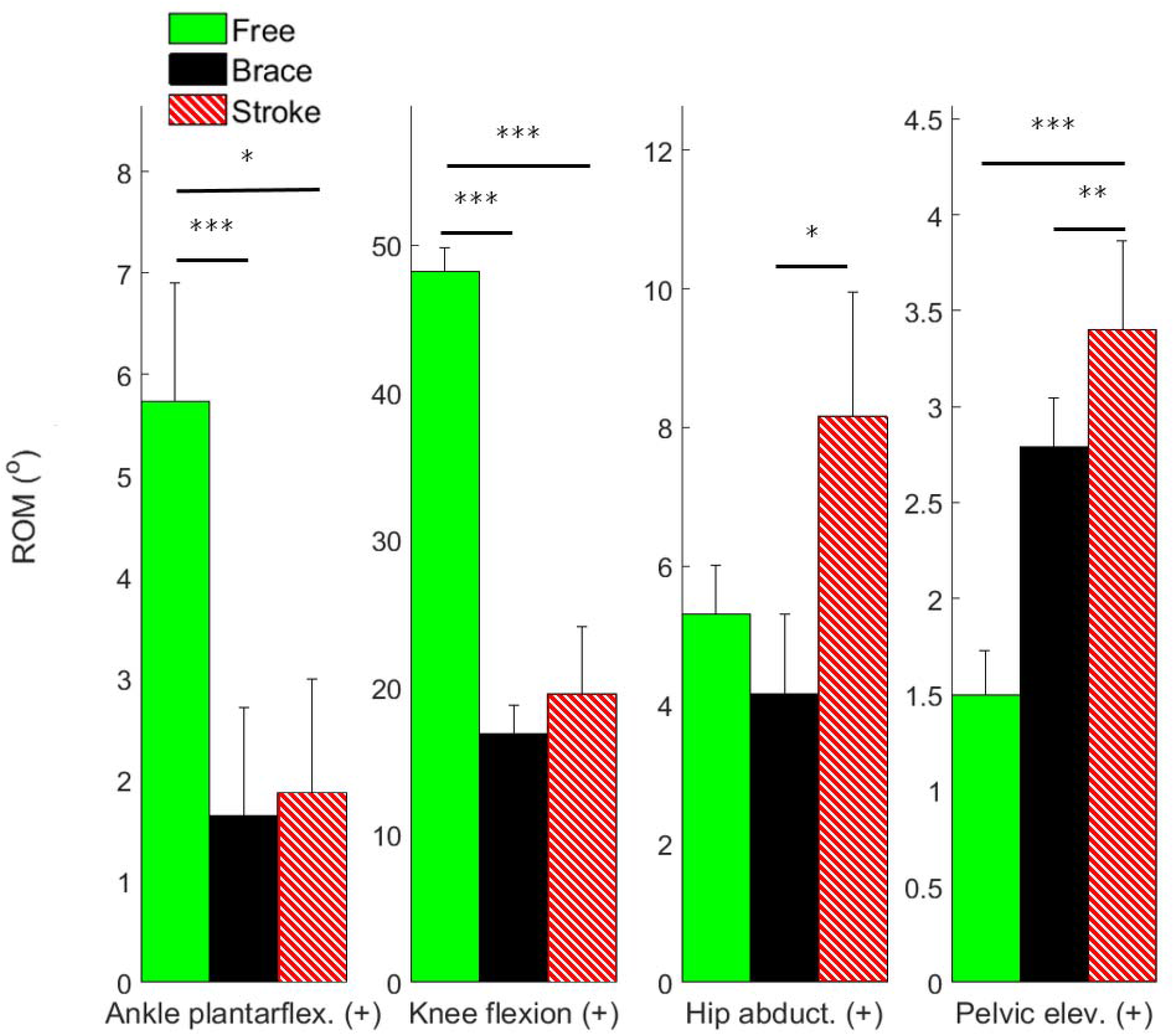
Similar responses in pelvic obliquity of *Brace* condition and post-stroke SKG but increased hip abduction in stroke gait. The bar graphs and the error bars represent the mean ROM values and standard errors respectively for knee flexion, ankle plantarflexion, hip abduction for constrained/paretic side and pelvic elevation in healthy individuals with *Free* and *Brace* conditions and stroke group. The level of significance between groups for given outcome measure were indicated on the lines connecting corresponding box plots (* p < 0.05, ** p < 0.01, *** p< 0.001).

**Table S1.**
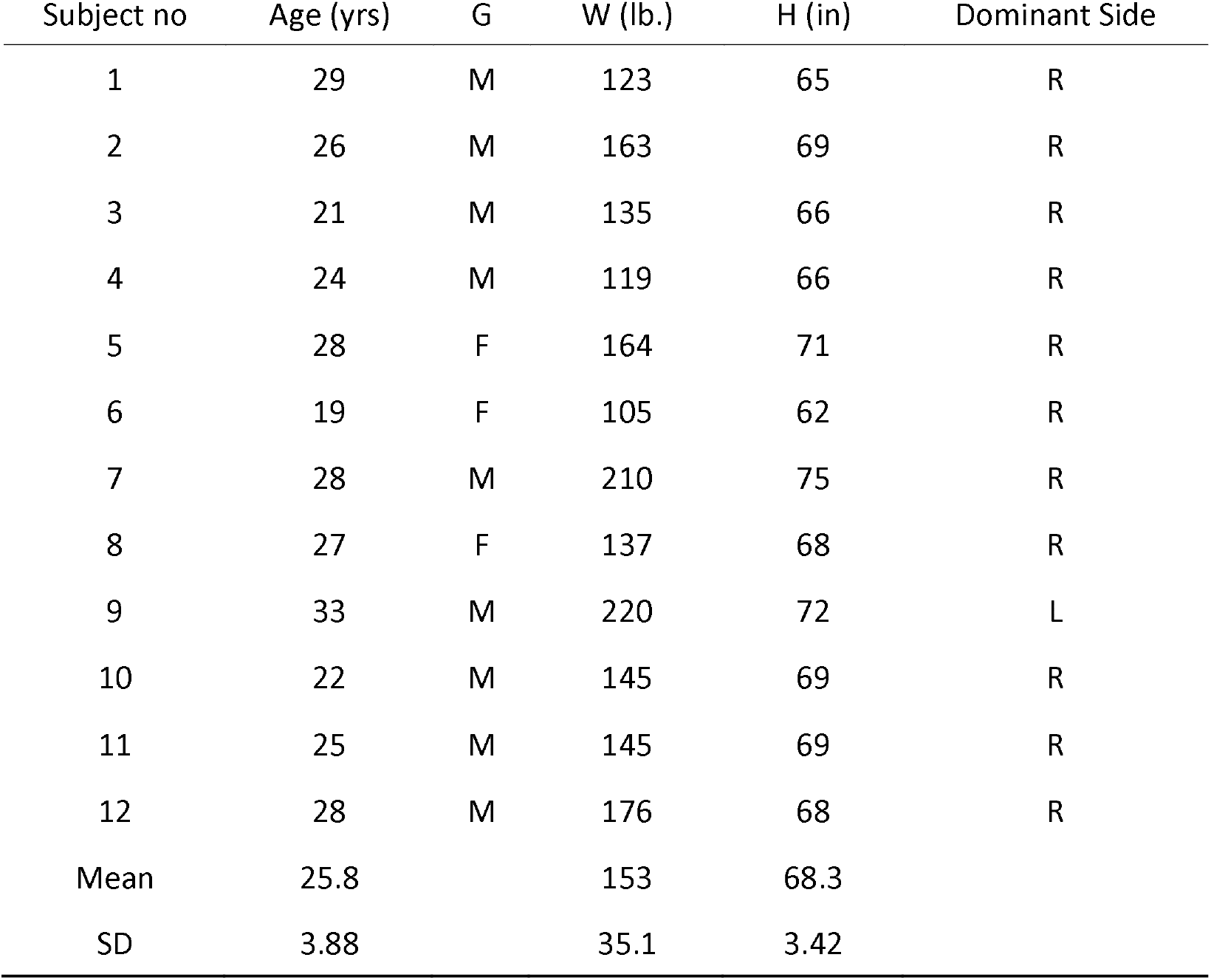
Demographics of healthy participants

**Table S2:** Summary of the joint angle ROM and spatiotemporal gait measures for healthy gait for Free, AFO, Brace and Restricted conditions with slow and normal walking speeds. The measures indicate the mean values and standard deviations. Due to the large complexity of markings, statistical comparisons are not noted on the table. Please see the text for the highlighted comparisons.

| Group<br>Outcome measure |  |  | Free | AFO | Brace | Restricted |
| --- | --- | --- | --- | --- | --- | --- |
| Kinematic measures | Knee ROM (°) | Slow | 48.2 ± 5.76 | 46.8 ± 6.86 | 16.9 ± 6.57 | 16.95 ± 7.09 |
|  |  | Normal | 56.1 ± 6.71 | 57.47 ± 7.10 | 21.73 ± 6.56 | 22.05 ± 6.94 |
|  | Hip abd. ROM(°) | Slow | 5.31 ± 2.46 | 5.46 ± 3.30 | 4.16 ± 3.98 | 4.24 ± 4.24 |
|  |  | Normal | 6.31 ± 4.24 | 5.08 ± 3.52 | 4.78 ± 3.21 | 4.86 ± 4.86 |
|  | Pel obl. ROM(°) | Slow | 1.50 ± 0.80 | 1.97 ± 0.89 | 2.79 ± 0.88 | 2.80 ± 0.94 |
|  |  | Normal | 1.40 ± 0.93 | 1.18 ± 0.84 | 2.82 ± 1.41 | 3.27 ± 1.49 |
|  | Ankle ROM(°) | Slow | 5.73 ± 4.04 | 3.72 ± 4.22 | 1.64 ± 3.72 | 1.61 ± 3.66 |
|  |  | Normal | 9.48 ± 6.83 | 6.26 ± 4.63 | 6.08 ± 5.07 | 3.47 ± 3.79 |
|  | Peak contralateral ankle plantarflexion (°) | Slow | 4.62 ± 3.40 | 4.61 ± 2.43 | 5.21 ± 3.65 | 3.75 ± 3.18 |
|  |  | Normal | 5.04 ± 3.31 | 4.92 ± 3.64 | 4.67 ± 2.75 | 5.68 ± 3.49 |
| Spatiotemporal measures | Max. toe height (mm) | Slow | 79.9 ± 42.1 | 65.8 ± 12.3 | 49.0 ± 17.6 | 54.7 ± 13.1 |
|  |  | Normal | 103 ± 42.2 | 86.3 ± 12.4 | 66.4 ± 16.9 | 62.8 ± 21.8 |
|  | Max. toe clearance (mm) | Slow | 9.41 ± 5.58 | 15.5 ± 5.73 | 10.9 ± 7.86 | 13.3 ± 4.26 |
|  |  | Normal | 13.82 ± 7.77 | 19.6 ± 8.86 | 17.0 ± 6.44 | 18.9 ± 12.6 |
|  | Max. toe width (mm) | Slow | 29.1 ± 9.12 | 29.3 ± 9.61 | 39.5 ± 11.5 | 38.5 ± 14.5 |
|  |  | Normal | 28.7 ± 10.2 | 32.5 ± 15.7 | 42.8 ± 20.0 | 44.1 ± 25.8 |
|  | Width at max. toe clearance (mm) | Slow | -8.82 ± 12.7 | -11.4 ± 14.8 | -15.9 ± 13.4 | -22.6 ± 20.4 |
|  |  | Normal | -7.09 ± 14.3 | -9.18 ± 25.1 | -26.3 ± 28.2 | -25.1 ± 45.1 |
|  | Swing ratio | Slow | 0.99 ± 0.05 | 1.07 ± 0.07 | 1.20 ± 0.14 | 1.20 ± 0.09 |
|  |  | Normal | 1.01 ± 0.04 | 1.02 ± 0.05 | 1.12 ± 0.17 | 1.21 ± 0.20 |
|  | Pre-swing ratio | Slow | 1.03 ± 0.09 | 0.93 ± 0.09 | 1.02 ± 0.10 | 0.97 ± 0.15 |
|  |  | Normal | 0.94 ± 0.06 | 0.89 ± 0.05 | 1.01 ± 0.12 | 0.98 ± 0.15 |

